# Noninvasive Laser SCOS Monitoring of Rat Brain Hemodynamics During Intracerebral Injection

**DOI:** 10.64898/2026.03.03.709425

**Authors:** Matthew Fernandes, Yu Xi Huang, Isabel Xu, Cesar Noguera Saigua, Junwei Li, Simon Mahler

**Affiliations:** Department of Biomedical Engineering, Stevens Institute of Technology, Hoboken, NJ 07030, USA; Department of Electrical Engineering, California Institute of Technology; Pasadena, CA 91125, USA; Glia Medical Inc., Irvine, CA 92620, USA

## Abstract

Cerebral blood volume (CBV) and blood flow (CBF) constitute key metrics for cerebrovascular monitoring, enabling assessment of stroke severity and risk-prediction, aging-related changes, and neurological diseases. CBF and CBV monitoring are key aspects in diagnosis, treatment triage, and clinical outcome of ischemic and hemorrhagic strokes. In recent years, there have been ongoing efforts toward the development of optical devices for noninvasive monitoring of CBV and CBF. Speckle contrast optical spectroscopy (SCOS) has recently emerged as a strong candidate for clinical translation in monitoring CBF and CBV, due to its affordability, compact and wearable design, and noninvasive nature. However, experimental demonstrations that SCOS can effectively monitor brain hemodynamics remain sparse. This is primarily due to challenges in design experiments that isolate cerebral blood dynamics from those in the scalp and skull.

In this paper, we report experiments using SCOS to monitor cerebral hemodynamics in rats during intracerebral blood flow modulation. To modify cerebral blood dynamics, a surgical procedure was performed to insert a catheter for direct injection of flow modulation fluids into the brain. Using the SCOS device, we monitored changes in CBV during deliberate CBF interventions into the brains of five rats. A saline solution was also injected as a sham control of the flow intervention. The results show a significant decrease in CBV during injection, followed by a return to baseline. This behavior is consistent with physiological expectations, as the injected fluids dilute the blood, leading to a transient reduction in blood volume. Notably, the CBV decrease induced by the flow modulation fluid solution required more than twice as long to recover to baseline compared with the saline solution, which is consistent with the delayed clearance of the flow modulation fluid by design. These experimental results demonstrate the effectiveness of SCOS for monitoring cerebral hemodynamics in animal models and highlight its potential for translation to human studies. Moreover, this work paves the way for the testing and characterization of cerebral therapeutic agents intended for blood flow modulation in animal models.

## Introduction

Cerebral blood volume (CBV) and cerebral blood flow (CBF) constitute important metrics for cerebrovascular monitoring and brain health assessment^1–3^. There have been ongoing efforts to develop transcranial optical devices capable of noninvasively monitoring CBV and CBF, providing real-time insights into brain perfusion.

Among these emerging techniques, speckle contrast optical spectroscopy (SCOS)^4–10^ (also called speckle visibility spectroscopy (SVS)^11–15^), has gained considerable attention to simultaneously measure CBV and CBF with high temporal resolution (> 60 Hz). Due to their noninvasive nature, affordability, compact and lightweight design, wearability, and ease of use, SCOS devices can be directly mounted on the head^4,5,16,17^, making them strong candidates for clinical translation in applications such as stroke ^4,10,16^ and brain injury assessment^17^.

In SCOS, CBV is estimated by quantifying optical signal attenuation^4,10^, while CBF is derived from speckle pattern analysis^6,7^. The ability to simultaneously provide noninvasive measurements of both CBV and CBF transcranial represents a significant advance toward real-time monitoring of cerebrovascular function.

Although SCOS shows exciting potential, its ability to accurately measure cerebral hemodynamics through the scalp and skull in both animals and humans has yet to be fully established^18^. While some experimental studies have demonstrated that SCOS can probe beyond the scalp and skull layers^11,18^, to the best of our knowledge, none have directly measured changes in brain hemodynamics alone using SCOS.

To further build on these, we report in this paper experiments using SCOS to monitor cerebral hemodynamics in rats during intracerebral blood flow intervention via fluid injection. Specifically, cerebral blood dynamics was intervened by an endovascular surgical procedure to directly inject flow modulation fluids into the cerebral arteries via a microcatheter.

Using SCOS, we tracked changes in CBV following injection of two types of mixtures into cerebral arteries: a saline solution (a sham control) and a flow modulation fluid (with intended hemodynamic modification effects), administered to five rats. The results revealed a significant decrease in CBV during injection, up to 12% with saline solution and up to 35% with the flow modulation fluid, followed by a return to baseline, consistent with physiological expectations that the injected solution transiently dilutes blood volume. Notably, we observe that the CBV decrease induced by the flow modulation fluid required more than twice as long to recover to baseline compared with saline.

These findings demonstrate the effectiveness of SCOS for monitoring cerebral hemodynamics in animal models and underscore its potential for translation to human studies. Furthermore, this work provides a foundation for future testing and characterization of therapeutic agents in CBF modulation and other cerebral solutions in preclinical models, advancing the development of tools for dynamic cerebrovascular monitoring, one of the key aspects in diagnosis of both ischemic and hemorrhagic strokes. CBV and CBF have been part of standard of care in ischemic stroke diagnosis, treatment triage and clinical outcome determination.

## Materials and Methods

### Speckle contrast optical spectroscopy (SCOS) for measuring cerebral blood volume (CBV)

In SCOS, a coherent laser beam illuminates the scalp and underlying tissues. As the light penetrates the tissue, it undergoes multiple scattering events. A scattering event occurs when incident photons are redirected from their original propagation direction without being absorbed, often accompanied by changes in their optical properties, such as direction, phase, or polarization. These multiple scattering events distort the coherent light beam, producing a granular interference pattern known as a speckle field (speckles). Speckles arise from the mutual interference of light traveling along different optical paths (i.e., accumulating different phase) within the scattering medium^19–21^, which can occur at rapid timescale (nanosecond)^22,23^. Fluctuations in the speckle field are sensitive to the underlying motion of components within the tissue, primarily red blood cells in our case and thus provide information about blood dynamics^15,19,24–26^. These temporal fluctuations in the speckle field are recorded using a high-pixel count (millions of pixels) and high-frame-rate count camera (60 frames per second or higher)^5,18,27^. From the recorded speckle patterns images, both blood volume and blood flow can be calculated.

### Cerebral blood volume (CBV) calculations

As mentioned earlier, motion of components within tissue induces temporal fluctuations in the speckle field, which in turn affect the total intensity of the detected speckle pattern. Because hemoglobin in red blood cells is a dominant absorber of near-infrared light, changes in hemoglobin concentration alter the amount of laser light collected back from the tissues. Consequently, the optical absorption coefficients can be related to the amount of hemoglobin content, thus the blood volume. By quantifying variations in the speckle field intensity and relating them to changes in optical attenuation, typically through Beer–Lambert law (e.g., Eq. 1), relative changes in cerebral blood volume (CBV) can be estimated as:

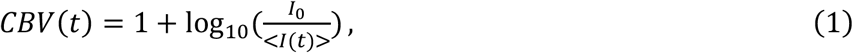

where I(t) is the intensity of the speckle field recorded by the camera, *I*_0_ is the intensity at baseline typically calculated as the average intensity of the entire image acquired during a baseline period of few seconds. The mean < *I*(*t*) > was calculated as the average spatial intensity of the entire image *I* acquired at time *t*, and can be expressed in camera grayscale units. The CBV in Eq. (1) was normalized to unity to improve visualization and facilitate identification of CBV reductions during injection.

It is important to note that our SCOS device provides a relative measurement of CBV, meaning that changes in CBV are reported relative to a baseline period within the same subject^17,18^. Relative CBV measurements reflect the fractional or percentage change from baseline, which is sufficient for monitoring dynamic responses, such as those induced by stimuli or interventions. In contrast, absolute CBV measurements provide the true blood volume unit (e.g. mL/100 g of brain), allowing comparisons across subjects or experimental conditions^28–30^. Achieving absolute CBV with optical methods is challenging because it requires accurate knowledge of tissue optical properties, light scattering, and absorption, which can vary across individuals and experimental setups. Therefore, SCOS is primarily used for tracking within-subject changes in CBV over time rather than reporting absolute volumetric values.

### Cerebral blood flow (CBF) calculations

While we primarily use CBV for the majority of the results presented in this paper (see the Results section for explanation), CBF can also be extracted from SCOS. In fact, CBF is the most commonly reported metric in SCOS studies.

In SCOS, CBF is obtained by quantifying the fluctuations in the speckle pattern over time, which result from the motion of red blood cells within the tissue. When blood flows quickly, the positions of red blood cells change rapidly within the camera exposure time. This rapid motion decorrelates the speckle pattern faster, washing out the speckles, i.e. reducing the standard deviation of intensity relative to its mean, which is what speckle contrast measures.

Contrarily, when blood flows slowly, the positions of red blood cells within the tissue change less during the camera exposure time. This slow motion decorrelates less the speckle pattern, enhancing its grainy pattern within the camera exposure time, i.e. increasing the standard deviation of intensity relative to its mean. If the blood is stationary (not flowing), the speckle pattern does not change over the camera exposure time, so the speckle contrast is maximal. In short, speckle contrast is inversely related to the blood speed: higher flow lowers speckle contrast and vice-versa. The speckle contrast is defined as the ratio of the standard deviation to the mean intensity of the recorded speckle patterns *I*(t):

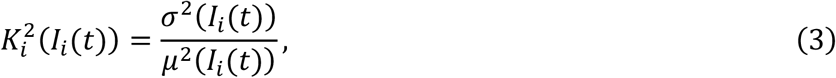

where *σ*^2^(*I*(*t*)) is the variance of the normalized image *I* of channel *i* at time *t* and *μ*(*I*(*t*)) its mean. The various sources of noise are accounted as ^5,6,31,32^:

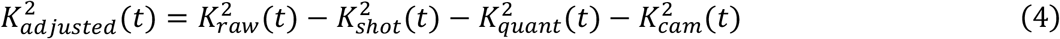

with 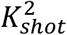 accounting for variance contributions from the shot noise, 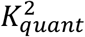 for the variance inherited from quantization, and 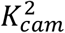 for the variance contributions of the camera’s readout noise and dark noise. See Ref. ^5^ for more details about the speckle contrast calculations and calibration processes. In implementing the K^2^-adjusted correction, all calibration and noise correction steps were performed in analog-to-digital units, i.e., grayscale counts. Camera readout noise was experimentally determined from the variance of dark frames, rather than relying on manufacturer specifications, to address potential CMOS degradation or inter-device variability. Shot noise, originally in electron units, is converted to analog-to-digital units. Quantization noise is minimal in our setup due to the use of 16-bit acquisition and was ignored in our measurements. These steps ensure that all noise sources are consistently expressed in the same units and appropriately scaled.

The cerebral blood flow (CBF) is related to 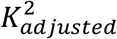 as:

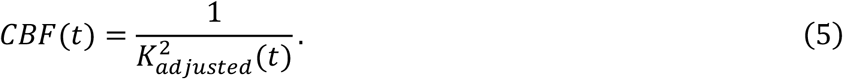

It is important to note that the definition relies on several key assumptions: (i) the camera’s exposure time is much longer than the speckle field’s decorrelation time, (ii) blood cell motion from multiple scattering events is random and unordered, and (iii) the scattered light field is ergodic with minimal contribution from static scatterers ^33^. Variations in scattering regimes and particle motion types can significantly influence the field correlation function *g*_1_(*τ*) ^33–35^, complicating the direct conversion of CBF to physical units (e.g., area/time) without further assumptions or corrections. In this study, we employed a min-max normalized CBF to highlight relative changes in blood flow rather than report absolute values.

### SCOS Experimental Arrangement

Figure 1 (a) shows a top-view of our SCOS experimental setup for measuring CBV and CBF in rats. The SCOS setup is fiber-based, consisting of a single illumination fiber that delivers near-infrared laser light to the rat’s head and a detection fiber positioned laterally on the other side of the rat’s head to collect the exiting light from the brain.

**Figure 1.**
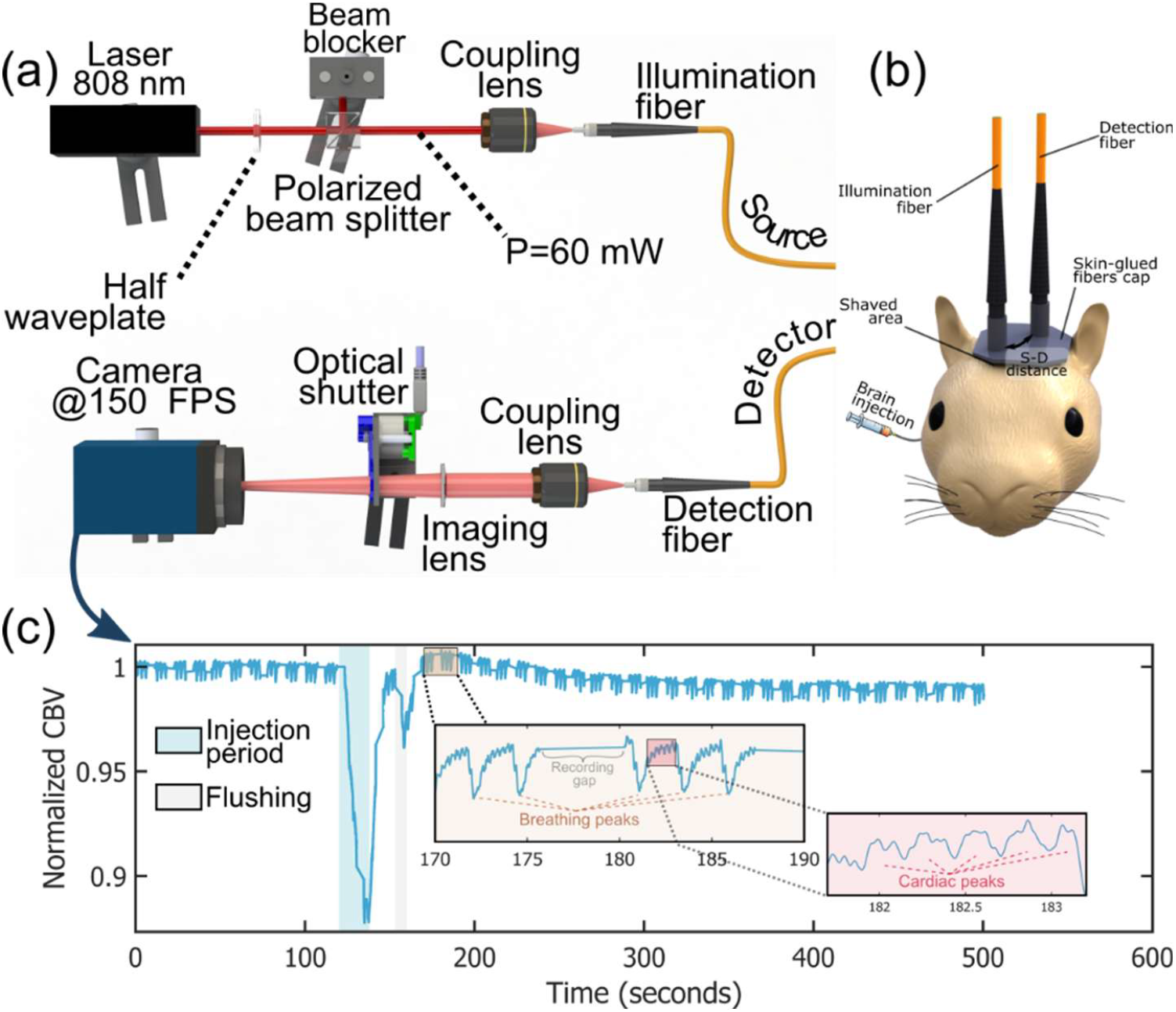
(a) Optical arrangement of the fiber SCOS device, consisting of a near-infrared laser source and a high-speed camera for speckle acquisition. (b) Illustration of the SCOS fibers placement on the rat’s head. One illumination fiber delivers laser light, and one detection fiber collects the backscattered light exiting the brain. A catheter is inserted into the right common carotid artery (CCA) while the right external carotid artery (ECA) was litigated to inject the mixture solution directly into the rat’s brain through right internal carotid artery (ICA). (c) Representative cerebral blood volume (CBV) time trace recorded during injection, showing a significant decrease in CBV during injection, demonstration the ability of the SCOS system to sensitively detect cerebral hemodynamics.

On the illumination part of the device, a single frequency infrared laser [Coherent ROUSB-808-PLR-110-1K] of 808 nm wavelength, with a large coherence length (> 1 m), and operating at 110 mW served as laser source. Although the laser source could provide up to 110 mW, the output power was limited to 60 mW for safety using a waveplate and a polarized beam splitter before coupling the laser light into the illumination optical fiber using a coupling lens. The illumination fiber was a 600 μm core diameter multimode fiber [Thorlabs M29L01] and was positioned few millimeters away from the rat’s scalp, producing a 5 mm illumination spot diameter, similarly to Refs.^5,10,11,17^. This setup ensured the laser intensity remained within the American National Standards Institute (ANSI) safety limit for skin exposure to an 808 nm laser beam (3.28 mW/mm^2^)^36^.

On the opposite side (detection fiber), a 600 μm core diameter multimode collected the exiting light from the rat’s head. The detection fiber was in direct contact with the skin to maximize light collection. The output of the fiber was imaged onto a scientific CMOS camera [Excelitas pco.edge 5.5] featuring a 6.5 × 6.5 μm pixel size and 16-bit pixel depth dynamic range using a coupling lens together with an imaging lens. The camera operated with an exposure time of 3 ms and was recording at around 150 FPS. This resulted in a speckle density of 0.1 speckle per pixel, corresponding to a one-dimensional speckle-to-pixel length ratio of s/p = 3^5,37,38^. To calibrate the various sources of noises (Equation 4), an optical shutter was placed in front of the camera.

Figure 1(b) illustrates the orientation and mounting of the optical fibers on the rat’s head. Prior to fiber placement, the scalp’s hair was removed using a depilatory shaving gel to minimize residual hair and improve optical coupling. The fibers were then inserted into a custom-made rubber cap that was glued to the rat’s scalp. The flexibility of the rubber cap allowed fine adjustment of fiber orientation while maintaining stable contact with the scalp throughout the experiment.

### Source-to-detector distance and brain sensitivity

A main component in transcranial optical measurements of CBF and CBV is the source-to-detector (S-D) distance, defined as the distance between the illumination point on the head (source) and the detection point (detector) where the scattered light is collected back, Figure 1(b).

As illumination light penetrates the scalp, it undergoes multiple scattering, with a fraction reaching the skull and brain before being scattered back toward the surface. A detection fiber placed at a source–detector (S– D) separation distance collects a small portion of this backscattered light. The depth to which the collected light penetrated into the head is intrinsically related to the S-D distance^39–41^, but the exact relationship between penetration depth and S-D distance remains unclear^11,18^. For a rat head model, typical anatomical layer thicknesses are approximately 1–2 mm for scalp, 0.5–2 mm for skull, and < 1 mm of cerebrospinal fluid and cortical tissue before reaching brain structures. This sets the skin to brain depth in rat head model to be less than 5 mm. Given these dimensions, a S-D distance of 1 to 1.5 cm is generally sufficient to probe brain signal in SCOS^11^. A study in small animal such as rabbit have demonstrated that S-D separations in this range balance adequate brain sensitivity with acceptable signal-to-noise ratio and manageable light attenuation for SCOS measurements^11^.

To minimize contamination from scalp blood flow, the detection fiber can be placed in direct contact with the scalp under slight mechanical pressure, effectively occluding superficial vasculature. This pressure-induced occlusion reduces the contribution of superficial blood dynamics to the measured signal, enhancing the relative sensitivity to cerebral blood flow.

In our setup, the source-detector distance between the illumination and detection fibers was comprised between 1 to 1.5 cm (typically around 1.2 cm). Note that in human studies, the typical S–D distance in compact SCOS devices ranges from approximately 3.5 to 4.5 cm^4,5,17^. In contrast, as mentioned earlier, for rodents and other small animals^11^, an S–D separation of 1 cm or greater is sufficient to achieve sensitivity to cerebral hemodynamics.

### Endovascular procedure and fluid injection to cerebral arteries

Adult Sprague-Dawley rats of 200-300 gram and around 16 weeks old were used for the animal study. Isoflurane was used as inhalant anesthetic during the procedure. An incision was made in the neck. Under microcopy, the common carotid artery (CCA), external carotid artery (ECA) and internal carotid artery (ICA) were surgically isolated around the bifurcation. A microcatheter (outer diameter of 0.5mm, and inner diameter around 0.2 mm) was inserted from the ECA to the CCA to deliver the treatment. Saline or flow modulation fluid was injected through the microcatheter into the CCA. The injected solution was then directed by the blood flow to the ICA and the cerebral circulation.

Around 0.2 ml of saline or flow modulation fluid or was injected by a 1 mL syringe via the microcatheter in the CCA. The duration of the injection was 30-60 seconds. The flow modulation fluid is supplied by Glia Medical Inc, Irvine CA, which has proprietary formulation that designed to slow down the local blood flow when injected cerebral vasculature and it also possessed a delayed clearance with blood circulation comparing with saline.

### Data recording protocol and challenges

For each rat, the following experimental protocol was applied:

1. The animal underwent a surgical procedure for catheter insertion and vascular bypass to enable intracerebral injection of the solution.
2. Following surgery, the hair on the scalp area above the brain was shaved.
3. The fiber-based SCOS device was positioned on the rat’s head. An initial baseline recording of 30 seconds was acquired to verify signal quality and system functionality.
4. The fiber cap was glued (using Gorilla super glue) to the scalp.
5. Once the fiber cap was glued, we started the recording of CBV and CBF as follows: a baseline period of 120 s was first recorded. At the 120 s time point, we started the intracerebral injection. To deliver the mixture, a 1 mL syringe was connected to the catheter (inner diameter: 0.2 mm) and prefilled with the solution prior to injection.
6. Immediately after the injection, the syringe containing the injected mixture was disconnected and replaced with a flushing syringe containing heparin saline of 5 units per mL to inject 0.1 mL. The flushing solution helps prevent blood coagulation within the catheter and preserves catheter patency for subsequent injections.
7. Following flushing, data acquisition was continued until 300 s for saline injections and up to 500 s for agent-solution injections to capture post-injection recovery dynamics.
8. Injection start and stop times were recorded manually.

We note the following challenges encountered during the recordings:

#### Challenge 1

Data acquisition began after a waiting period of 5 to 10 min to allow partial drying of the glue. Because the surgical procedure limited the viable recording time to only a few hours per animal, the glue was not fully dry (typically requires 60 min). As discussed later, adhesive fixation proved suboptimal as a mounting strategy, and improved attachment methods will be explored in future studies (see Discussion).

#### Challenge 2

Accurate placement of the fiber cap was critical, as the margin for error was 1 to 2 mm. Indeed, given the source–detector separation of 1 to 1.5 cm, and the small accessible brain surface flat area in rats, a misplacement on the order of 1 to 2 mm could result in lower brain sensitivity. Consequently, in some animals, slight misalignment of the device likely reduced brain sensitivity and contributed to lower data quality (see Results). Incorporating an adjustable head strap for device mounting would improve SCOS placement accuracy and stability (see Discussion).

#### Challenge 3

The rats were not positioned in a stereotaxic frame or rigid head-holding apparatus during the experiments, which we acknowledge as a limitation of the current study. Although the optical fibers were secured (via tape) to external supports (pole and table) to reduce motion, residual head movement caused mechanical perturbations of the fibers and device. As a result, motion artifacts, particularly those associated with breathing, were apparent in the recorded signals.

While CBV measurements remained sufficiently robust under these conditions, CBF, which is more sensitive to motion, could not be reliably extracted. Accordingly, the results presented in this paper focus exclusively on CBV, despite SCOS being capable of measuring both CBV and CBF.

Future iterations of the experimental setup will incorporate stereotaxic head fixation and improved device mounting to minimize motion artifacts and ensure precise positioning. These improvements are expected to enable simultaneous, reliable measurements of both CBV and CBF (see Discussion).

#### Challenge 4

The injection durations varied across trials and depended on the injected mixture as well as external factors, including manual injection speed and the quality of the catheter connection. In future studies, a syringe pump will be used to standardize injection rate and duration (see Discussion). Similarly, because flushing was performed manually, its duration varied between realizations, representing another source of variability that will be addressed in future experimental iterations (see Discussion).

#### Challenge 5

The communication speed and computational capabilities were insufficient to support continuous CBV and CBF recording over the full 300 s or 500 s duration. As a result, data acquisition was segmented into repeated cycles consisting of 8 s of recording followed by a 2 s pause, during which the recorded data were processed to extract CBV and CBF before acquisition resumed. This limitation is primarily hardware-related and can be addressed in future implementations by using a higher-performance CPU and GPU. The computer used in this study was equipped with CPU: Intel Core™ i7-11700K, GPU: NVIDIA GeForce GTX 1660 Ti, 6 GB GDDR6.

In the Discussion section, we propose alternative approaches and solutions to address these challenges, which we plan to implement in future studies. This work serves as a proof-of-concept demonstrating the effectiveness of SCOS for monitoring cerebral hemodynamics in animal models.

## Results

Figure 1(c) shows a representative CBV time trace recorded during injection of 0.2 mL of saline solution. As evident, CBV decreases significantly following the injection onset at 120 s and rapidly returns to baseline once the injection ends. The CBV decrease observed at injection is consistent with physiological expectations. During injection, the saline solution enters the cerebral vasculature and dilutes the blood. As blood becomes more diluted, its optical absorption decreases because hemoglobin is the dominant absorber of near-infrared light. Consequently, the reduced absorption leads to a decrease in the measured CBV, as described by Equation (1). The effect of the flushing injection on CBV is also visible.

Because the injection is delivered exclusively into the cerebral compartment (i.e. the brain) and does not directly affect the scalp or skull layers, the observed CBV changes can be attributed to cerebral rather than extracerebral hemodynamic effects. This result provides experimental evidence that the SCOS system is sensitive to brain-specific blood dynamics in rats, despite signal contributions from overlying tissues. The ability to detect a localized, intracerebral perturbation further supports the depth sensitivity of SCOS and its capability to probe cerebral hemodynamics through the scalp and skull, which remains a central challenge for noninvasive optical techniques.

As shown in the insets of Figure 1(c), periodic fluctuations corresponding to the animal’s breathing and cardiac activity become evident upon zooming into the signal. These features enable extraction of the breathing rate (BR) and heart rate (HR) during injection using Fourier-based peak detection, as previously demonstrated in Refs.^11,18^.

Next, the experiment shown in Figure 1(c) was repeated across five different rats, with the corresponding CBV time traces presented in Figure 2(a). As shown, inter-animal variability is observed in the magnitude of the CBV decrease. A pronounced CBV drop at injection period is evident for rats #1 and #4, moderate for rat #3, and a smaller decrease for rats #2 and #5. Because the injected solution was saline and therefore not expected to induce chemical or pharmacological effects, these differences cannot be attributed to the mixture itself. Instead, we attribute the observed variability primarily to differences in device placement on the rats’ heads (see Challenge 2 in Materials and Methods).

**Figure 2.**
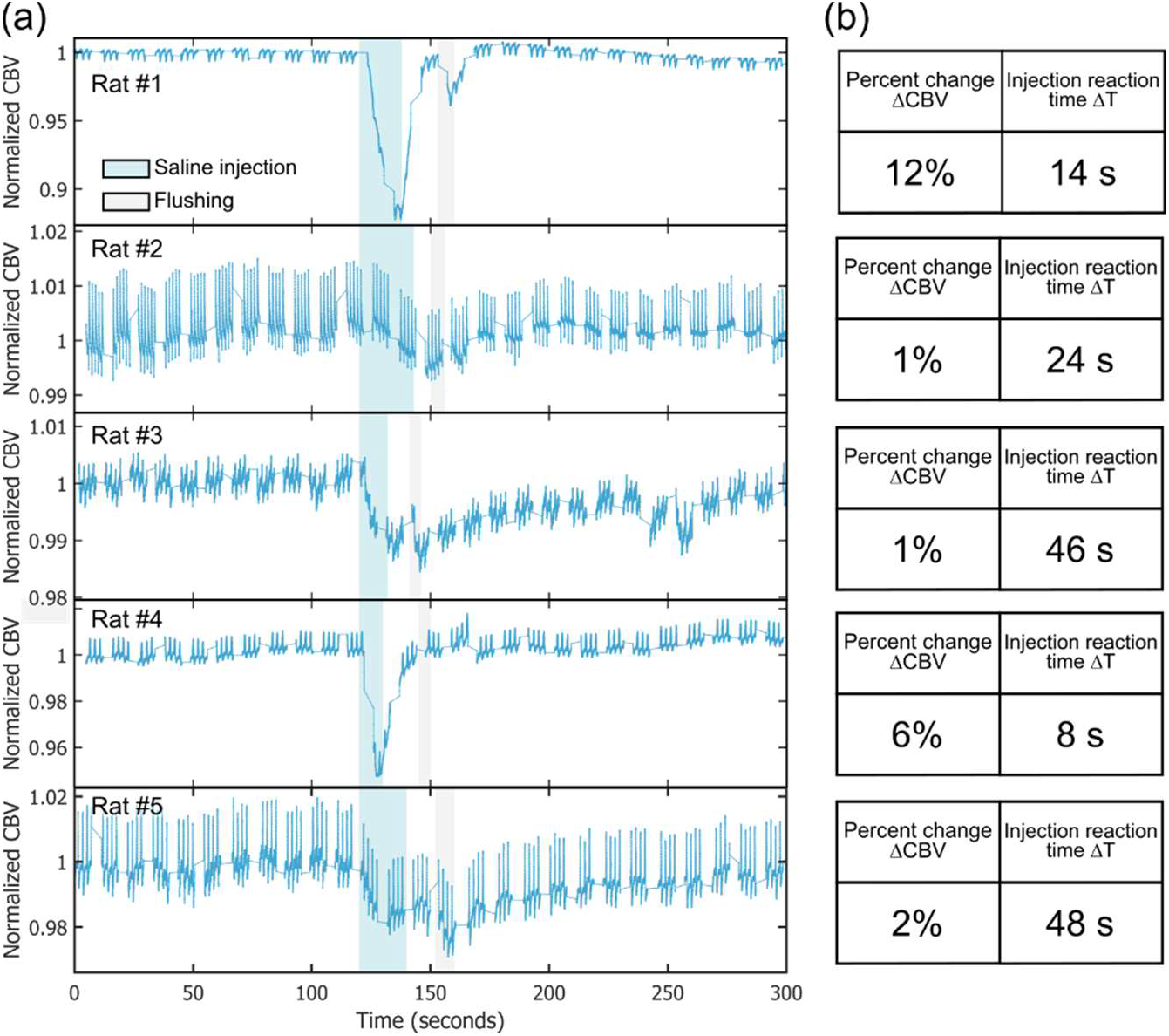
Saline solution intracerebral injection in rats. (a) Cerebral blood volume (CBV) time trace during saline injection from five different rats. (b) Extracted percent change of ΔCBV and injection reaction time ΔT using a Gaussian fit.

Accurate positioning of the fiber cap was critical, as the allowable margin for error was on the order of 1– 2 mm, a condition that was not consistently met across all animals. As a result, when the SCOS system is mispositioned, the measurements are more strongly influenced by scalp and skull signals, leading to a reduced apparent CBV decrease during injection. These observations further indicate that the CBV drop cannot be observed by measuring scalp-only CBV, reinforcing the cerebral origin of the detected signal.

While this limitation is unfortunate and will be addressed in future studies, the present work serves as a proof-of-concept. Importantly, all five rats exhibited a detectable CBV decrease during injection, supporting the robustness and validity of the SCOS measurements. We note that for all five rats, the written down injection times correlate well with the observed drop of CBV.

Finally, as noted earlier (Challenge 3), the rats were not positioned in a head-holding apparatus during the experiments, and residual head motion introduced mechanical perturbations to the fibers and device. As a result, motion artifacts, particularly affecting CBF measurements, corrupted the signal. Consequently, the results presented in this paper focus exclusively on CBV, despite SCOS being capable of measuring both CBV and CBF.

Next, we applied a Gaussian fit to the CBV time trace using the Gaussian function G(t):

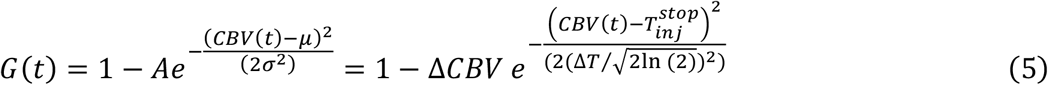

With A the amplitude of the Gaussian, corresponding to the percent change in CBV (*A* = Δ*CBV*), *μ* represents the center (mean) of the Gaussian peak and corresponds to the time at which injection stops 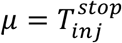, and *σ* is the Gaussian standard deviation. The standard deviation *σ* is related to the full width at half maximum (FWHM) of the Gaussian by 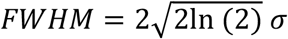. The injection reaction time Δ*T* is defined as half of the FWHM, i.e. 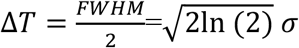.

The Gaussian fit was used to characterize the mechanical component of the CBV response during intracerebral injection of a transparent liquid (e.g., saline or agent solution). Under the assumption that the dominant effect is transient mechanical blood displacement, the response is expected to be temporally symmetric, with the decrease during injection mirroring the return to baseline. In this context, a Gaussian provides a reasonable first-order approximation of the mechanical transient. However, when chemical or biological effects contribute to the signal (particularly for the agent solution in Fig. 4), the response may become asymmetric, with distinct injection-driven and recovery dynamics. In such cases, a two-component fit (e.g., dual power-law or biexponential functions) would more appropriately decouple these processes and better capture the underlying physiology.

From the Gaussian fit, we extracted the percent change of Δ*CBV* and the injection reaction time Δ*T*. As shown in Figure 2(b), rats #1 and #4 exhibited the largest percent changes in CBV, 12% and 5%, respectively. In contrast, the injection reaction time (ΔT) was shortest for rats #1 and #4, measuring 14 s and 8 s, respectively. All saline injection reaction times were below 48 s.

Next, a Fourier transform was applied to each of the 8 seconds segments in the CBV time traces. From the Fourier frequencies, we identified and extracted the breathing rate (BR) and heart rate (HR)^11^. The resulting data were then categorized into three periods: before, during, and after injection. We note that across all rats, heart rate oscillated between 200 and 300 bpm, while breathing rate varied between 10 and 100 bpm.

The results are shown in Figure 3(a) for heart rate and Figure 3(b) for breathing rate. As observed, there were no consistent changes in either heart rate or breathing rate during or after saline injection compared with the baseline period. The largest heart rate change measured was less than 8%, with most variations falling within the measurement error (*ΔHR* = ±8 bpm). These results therefore indicate that the saline injection had no detectable effect on the animal’s heart rate or breathing rate. We also analyzed the cardiac pulse waveform in the CBV time traces before, during, and after injection for each rat and found no consistent changes in waveform pattern, indicating that the saline injection did not affect cardiac dynamics.

**Figure 3.**
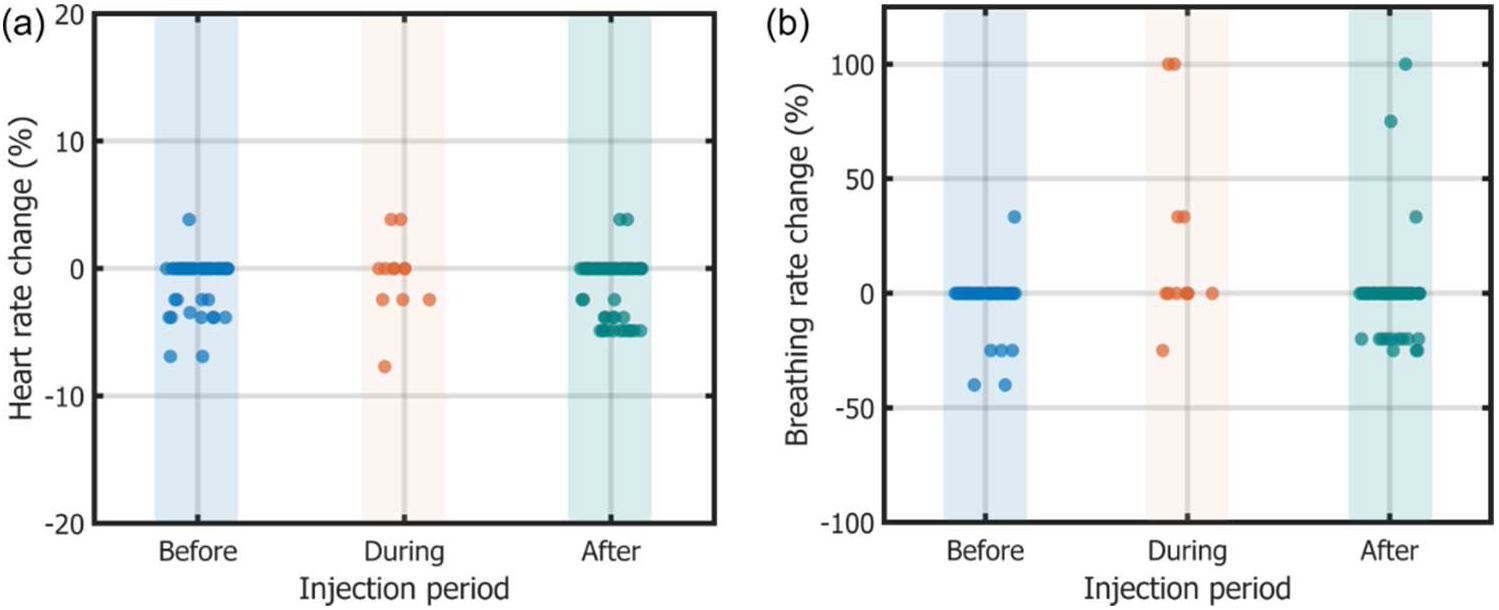
(a) Heart rate change and (b) breathing rate change for each injection period for all rats of Figure. 2.

Next, we repeated the experiments shown in Figure 2, this time injecting the agent solution instead of saline. This solution contained chemical compounds capable of affecting blood circulation. As our goal was not to characterize the effects of a specific drug but rather to demonstrate the capability of the SCOS system to monitor such changes, we did not control for the chemical composition or concentration of the agent in the mixture. For this experiment, we report results from six rats. As shown in Figure 4(a), the CBV responses to the agent solution were generally more pronounced than those observed with saline. A substantial CBV decrease during the injection period is evident for rats #1, #4, #5, and #6, while rats #2 and #3 exhibited moderate decreases. The largest observed percent change in CBV (ΔCBV) was 35% (Figure 4(b)).

**Figure 4.**
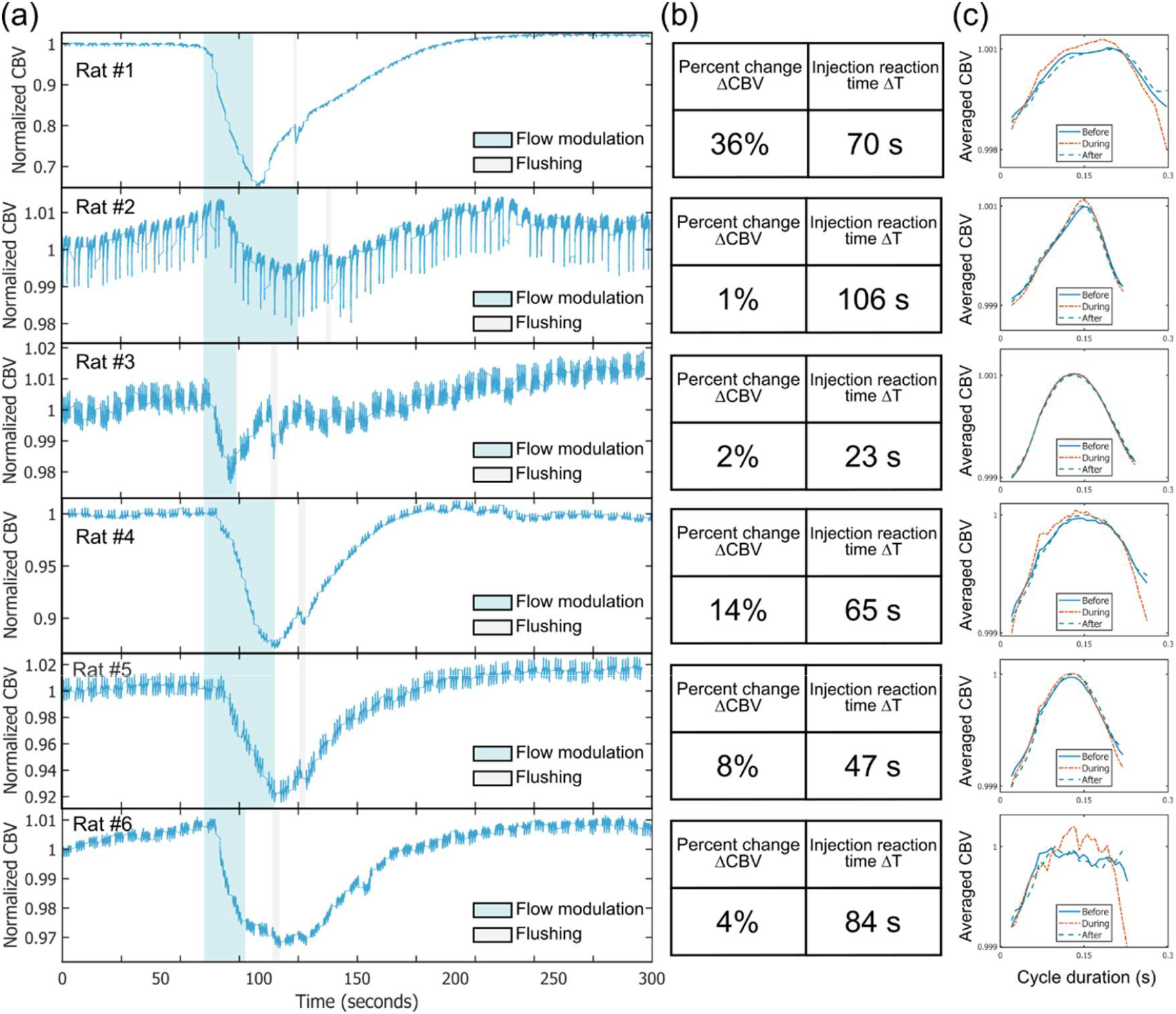
Flow modulation fluid intracerebral injection in rats. Cerebral blood volume (CBV) dynamics, measured in units of blood volume index (BVI), over a 5,240-second period with multiple injection events indicated as green areas.

The average percent change in CBV for the agent solution was 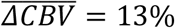 and the average injection reaction time was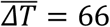. In comparison, for the saline solution (Figure 2(b)), 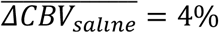 and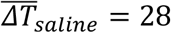, corresponding to roughly three times the CBV change and twice the injection reaction time for the agent solution. Even when comparing the best-case example (Rat #1 in Figures 2 and 4), the agent solution produced approximately three times greater CBV change and five times longer reaction time. These results indicate that the effects of the agent solution extend beyond the purely mechanical effects of blood dilution as with the saline solution. Thus, the observed increase in CBV percent change and injection reaction time can be attributed to the chemical compounds of the mixture. Overall, these findings demonstrate the potential of SCOS devices for evaluating and characterizing cerebral therapeutic agents in animal models.

Figure 4(c) shows the averaged CBV cardiac pulse waveform for each rat before, during, and after injection. As shown, no consistent changes in waveform pattern can be observed, indicating that the agent solution did not affect cardiac dynamics, as observed with the saline solution.

## Discussions

Overall, these experimental results demonstrate that SCOS is effective for monitoring CBV changes in animal models and is sensitive to both mechanical and chemical perturbations of the brain vasculature. This study serves as a proof-of-concept, highlighting the potential of SCOS for preclinical evaluation of cerebral therapeutic agents and paving the way for future translation to human studies.

Future improvements in device mounting, head stabilization, and injection control will further enhance the ability of SCOS to simultaneously measure both CBV and CBF, enabling more comprehensive characterization of cerebral hemodynamics. Specifically, we plan on:

– Custom 3D-printed head mount: a strap-on, adjustable mount will provide better mechanical stability while allowing the device to slide across the rat’s head to maximize brain signal. This will enable precise placement of the optical fibers.
– Stereotaxic positioning: placing the rats in a stereotaxic frame will minimize head and respiratory movements, reducing motion artifacts and enabling simultaneous measurement of CBF and CBV. The rat’s head can be secured using ear bars and a bite bar.
– Automated injection system: using a syringe pump will standardize injection duration and volume, improving experimental reproducibility and reducing variability caused by manual injections.
– Enhanced computational and acquisition capabilities: upgrading to a computer with at least a 16 GB GPU (e.g. NVIDIA RTX 4060 Ti 16 GB) and replacing the PCO camera (requiring a dedicated PCIe card) with a compact USB-based camera (e.g., Basler daA1920-160um^5^) should allow enough computational power for continuous recording of CBF and CBV over a 300 s period.
– Improved light collection: using a larger-diameter collection fiber (e.g., 1 mm instead of 600 μm) will increase the collected signal and could improve data quality.

With these improvements, we anticipate obtaining reproducible CBV and CBF responses across animals, enabling detailed characterization of the differences between saline and others agent solutions injections with statistical significance.

## Conclusion

In this study, we used a SCOS device to monitor cerebral hemodynamics in rats during intracerebral injection of two types of solutions: saline and the flow modulation fluid. To isolate cerebral blood dynamics, a surgical procedure was performed in which the right common carotid artery was bypassed and a catheter was inserted to allow direct injection. Using the SCOS device, we measured changes in cerebral blood volume (CBV) in five rats injected with saline and six rats injected with the flow modulation fluid.

For the saline injections, a pronounced CBV decrease during the injection period was observed in two rats, a moderate decrease in one rat, and a small decrease in the remaining two rats. In contrast, for the flow modulation fluid, four rats exhibited a substantial CBV decrease during injection, while two rats showed moderate decreases. The largest observed percent change in CBV (ΔCBV) was 35% (Figure 4(b)).

On average, the percent change in CBV for the saline solution was 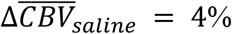, with an average injection reaction time of 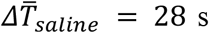. In comparison, for the agent solution, 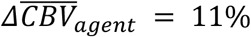 and 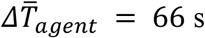, corresponding to approximately three times the CBV change and twice the injection reaction time for the agent solution. The flow modulation fluid had a proprietary formulation that designed to slow down the local blood flow when injected cerebral vasculature and it also possessed a delayed clearance with blood circulation comparing with saline. These results indicate that the effects of the agent solution extend beyond simple mechanical blood dilution, reflecting the action of chemical compounds on cerebral hemodynamics.

## Acknowledgement

The authors thanks professor Changhuei Yang for their support and help in the conceptualization of this project.

## Ethical considerations

All animal protocols were performed in conformity with the Animal Research: Acculab Life Sciences guidelines. Experimental procedures were approved by the Acculab Life Science animal care and use committees (IACUC) under protocol #219 and were performed under relevant personal and project licenses. The animal rat study was conducted at Acculab Life Sciences, a credited commercial lab with a PHS Approved Animal Welfare Assurance D16-00907 (A4716-01). Rat model was selected due to its wide use in literature reports for pre-clinical stroke models, artery size for endovascular access, and low cost.

## References

1. Rahul, J. S. & Kakkar, G. Cerebral Blood Flow Monitoring. in Principles and Practice of Neurocritical Care (eds Prabhakar, H., Singhal, V., Zirpe, K. G. & Sapra, H.) 75–92 (Springer Nature Singapore, Singapore, 2024). doi:10.1007/978-981-99-8059-8_6.

2. Yoshida, K. et al. Dynamics of cerebral blood flow and metabolism in patients with cranioplasty as evaluated by 133Xe CT and 31P magnetic resonance spectroscopy. Journal of Neurology, Neurosurgery & Psychiatry 61, 166–171 (1996).

3. Powers, W. J. Cerebral hemodynamics in ischemic cerebrovascular disease. Annals of Neurology 29, 231–240 (1991).

4. Favilla, C. G. et al. Validation of the Openwater wearable optical system: cerebral hemodynamic monitoring during a breath-hold maneuver. Neurophoton. 11, (2024).

5. Huang, Y. X. et al. Compact and cost-effective laser-powered speckle contrast optical spectroscopy fiber-free device for measuring cerebral blood flow. J. Biomed. Opt. 29, 067001 (2024).

6. Kim, B. et al. Measuring human cerebral blood flow and brain function with fiber-based speckle contrast optical spectroscopy system. Commun Biol 6, 844 (2023).

7. Valdes, C. P. et al. Speckle contrast optical spectroscopy, a non-invasive, diffuse optical method for measuring microvascular blood flow in tissue. Biomed. Opt. Express 5, 2769 (2014).

8. Dragojević, T. et al. Compact, multi-exposure speckle contrast optical spectroscopy (SCOS) device for measuring deep tissue blood flow. Biomed. Opt. Express 9, 322 (2018).

9. Lin, C.-H. P. et al. Multi-mode fiber-based speckle contrast optical spectroscopy: analysis of speckle statistics. Opt. Lett. 48, 1427 (2023).

10. Huang, Y. X. et al. Correlating stroke risk with non-invasive cerebrovascular perfusion dynamics using a portable speckle contrast optical spectroscopy laser device. Biomedical Optics Express 15, 6083–6097 (2024).

11. Mahler, S. et al. Assessing depth sensitivity in laser interferometry speckle visibility spectroscopy (iSVS) through source-to-detector distance variation and cerebral blood flow monitoring in humans and rabbits. Biomed. Opt. Express 14, 4964 (2023).

12. Huang, Y. X., Mahler, S., Mertz, J. & Yang, C. Interferometric speckle visibility spectroscopy (iSVS) for measuring decorrelation time and dynamics of moving samples with enhanced signal-to-noise ratio and relaxed reference requirements. Opt. Express, OE 31, 31253–31266 (2023).

13. Xu, J., Jahromi, A. K., Brake, J., Robinson, J. E. & Yang, C. Interferometric speckle visibility spectroscopy (ISVS) for human cerebral blood flow monitoring. APL Photonics 5, 126102 (2020).

14. Bandyopadhyay, R., Gittings, A. S., Suh, S. S., Dixon, P. K. & Durian, D. J. Speckle-visibility spectroscopy: A tool to study time-varying dynamics. Review of Scientific Instruments 76, 093110 (2005).

15. Dong, Z. et al. Non-invasive laser speckle contrast imaging (LSCI) of extra-embryonic blood vessels in intact avian eggs at early developmental stages. Biomed. Opt. Express 15, 4605 (2024).

16. Favilla, C. G. et al. Portable cerebral blood flow monitor to detect large vessel occlusion in patients with suspected stroke. J NeuroIntervent Surg jnis-2024-021536 (2024) doi:10.1136/jnis-2024-021536.

17. Mahler, S. et al. Portable six-channel laser speckle system for simultaneous measurement of cerebral blood flow and volume with potential applications in characterization of brain injury. Neurophoton. 12, (2025).

18. Huang, Y. X. et al. Assessing human scalp and brain blood flow sensitivities via superficial temporal artery occlusion using speckle contrast optical spectroscopy. APL Bioengineering 9, 046106 (2025).

19. Boas, D. A. & Dunn, A. K. Laser speckle contrast imaging in biomedical optics. J. Biomed. Opt. 15, 011109 (2010).

20. Senarathna, J., Rege, A., Li, N. & Thakor, N. V. Laser Speckle Contrast Imaging: theory, instrumentation and applications. IEEE Rev Biomed Eng 6, 99–110 (2013).

21. Chriki, R., Mahler, S., Tradonsky, C., Friesem, A. A. & Davidson, N. Real-time full-field imaging through scattering media by all-optical feedback. Phys. Rev. A 105, 033527 (2022).

22. Chriki, R. et al. Spatiotemporal supermodes: Rapid reduction of spatial coherence in highly multimode lasers. Phys. Rev. A 98, 023812 (2018).

23. Mahler, S. et al. Fast laser speckle suppression with an intracavity diffuser. Nanophotonics 10, 129–136 (2020).

24. Readhead, C. et al. Automated non-invasive laser speckle imaging of the chick heart rate and extraembryonic blood vessels and their response to Nifedipine and Amlodipine drugs. Developmental Biology 519, 46–54 (2025).

25. Durduran, T. et al. Diffuse optical measurement of blood flow, blood oxygenation, and metabolism in a human brain during sensorimotor cortex activation. Opt. Lett. 29, 1766 (2004).

26. Draijer, M., Hondebrink, E., Van Leeuwen, T. & Steenbergen, W. Review of laser speckle contrast techniques for visualizing tissue perfusion. Lasers Med Sci 24, 639–651 (2009).

27. Cheng, T. Y. et al. Choosing a camera and optimizing system parameters for speckle contrast optical spectroscopy. Sci Rep 14, 11915 (2024).

28. Uh, J., Lewis-Amezcua, K., Varghese, R. & Lu, H. On the measurement of absolute cerebral blood volume (CBV) using vascular-space-occupancy (VASO) MRI. Magnetic Resonance in Med 61, 659–667 (2009).

29. Muizelaar, J. P., Fatouros, P. P. & Schröder, M. L. A New Method for Quantitative Regional Cerebral Blood Volume Measurements Using Computed Tomography. Stroke 28, 1998–2005 (1997).

30. Engvall, C. et al. Human Cerebral Blood Volume (CBV) Measured by Dynamic Susceptibility Contrast MRI and 99mTc-RBC SPECT. Journal of Neurosurgical Anesthesiology 20, 41–44 (2008).

31. Sheppard, W. F. On the Calculation of the most Probable Values of Frequency-Constants, for Data arranged according to Equidistant Division of a Scale. Proceedings of the London Mathematical Society s1-29, 353–380 (1897).

32. Kobayashi Frisk, L. et al. Comprehensive workflow and its validation for simulating diffuse speckle statistics for optical blood flow measurements. Biomed. Opt. Express 15, 875 (2024).

33. Liu, C., Kılıç, K., Erdener, S. E., Boas, D. A. & Postnov, D. D. Choosing a model for laser speckle contrast imaging. Biomed. Opt. Express 12, 3571 (2021).

34. Boas, D. A. et al. Establishing the diffuse correlation spectroscopy signal relationship with blood flow. Neurophotonics 3, 031412 (2016).

35. Postnov, D. D., Tang, J., Erdener, S. E., Kılıç, K. & Boas, D. A. Dynamic light scattering imaging. Sci. Adv. 6, eabc4628 (2020).

36. American National Standard for Safe Use of Lasers, ANSI Z136.1 – 2022. (Laser Institute of America, Orlando, FL, 2022).

37. Robinson, M. B. et al. Comparing the performance potential of speckle contrast optical spectroscopy and diffuse correlation spectroscopy for cerebral blood flow monitoring using Monte Carlo simulations in realistic head geometries. Neurophoton. 11, (2024).

38. Zilpelwar, S. et al. Model of dynamic speckle evolution for evaluating laser speckle contrast measurements of tissue dynamics. Biomed. Opt. Express 13, 6533 (2022).

39. Strangman, G. E., Li, Z. & Zhang, Q. Depth Sensitivity and Source-Detector Separations for Near Infrared Spectroscopy Based on the Colin27 Brain Template. PLoS ONE 8, e66319 (2013).

40. Haeussinger, F. B. et al. Simulation of Near-Infrared Light Absorption Considering Individual Head and Prefrontal Cortex Anatomy: Implications for Optical Neuroimaging. PLoS ONE 6, e26377 (2011).

41. Myllylä, T. et al. Assessment of the dynamics of human glymphatic system by near-infrared spectroscopy. Journal of Biophotonics 11, e201700123 (2018).

